# Anterograde Trafficking of Toll-Like Receptors Requires the Cargo Sorting Adaptors TMED-2 and 7 and specific N-glycans

**DOI:** 10.1101/2022.10.10.511598

**Authors:** Julia E.J. Holm, Sandro G. Soares, Martyn F. Symmons, Martin C. Moncrieffe, Nicholas J. Gay

## Abstract

Toll-Like Receptors (TLRs) play a pivotal role in immunity by recognizing conserved structural features of pathogens and initiating the innate immune response. TLR signaling is subject to complex regulation that remains poorly understood. Here we show that two small type I transmembrane receptors, TMED2 and 7, that function as cargo sorting adaptors in the early secretory pathway are required for transport of TLRs from the ER to Golgi. Protein interaction studies reveal that TMED7 interacts with TLR2, TLR4, and TLR5 but not with TLR3 and TLR9. On the other hand, TMED2 interacts with TLR2, TLR4 and TLR3. Dominant negative forms of TMED7 suppress the export of cell surface TLRs from the ER to the Golgi. By contrast TMED2 is required for the ER-export of both plasma membrane and endosomal TLRs. We also find that a specific N-linked glycan of TLR2 is required for anterograde trafficking and may be directly recognised by TMEDs. Together, these findings suggest that association of TMED2 and TMED7 with TLRs facilitates anterograde transport from the ER to the Golgi.

## INTRODUCTION

Due to the crucial role of TLRs in immune responses, their signalling activity must be tightly regulated to avoid signalling dysfunction. Hence, TLRs are subjected to multiple layers of regulation at the levels of ligand activation, signal transduction, biosynthesis and trafficking. Toll-like receptors are translated into the endoplasmic reticulum (ER), where they are folded and subjected to post-translational modifications such as N-linked glycosylations and the formation of disulphide bonds (1, 2). Properly folded and processed proteins are incorporated into vesicles and transported to the Golgi compartments. In the Golgi they are subjected to further post-translational modifications. Mature TLRs are then sorted and packaged into vesicles directed towards their respective destination. However, this is only a general description of the biosynthetic pathway for glycoproteins such as TLRs and many questions remain unanswered regarding the particular mechanisms behind the transport of TLRs and the role of the biosynthesis in the regulation of TLR signalling.

In the ER, protein folding is assisted by various chaperone proteins and a few of these are involved in protein folding, sorting and transport of TLRs. Endosomal TLRs (TLR3, 7, 8 and 9) are associated with UNC93B1 in the endoplasmic reticulum (ER), a chaperone that mediates their trafficking to endolysosomes (3). Regarding TLR4, several proteins have been shown to associate with the receptor in the secretory pathway although their exact roles are not fully understood. For example, the ER chaperone gp96 (HSP90β1) is highly conserved in eukaryotes and is known to promote correct protein folding (4). Chaperone PRAT4A (protein-associated with TLR4) associates with TLR4 in the later phase of the ER maturation and is believed to recognize correctly folded and glycosylated TLR4 (5, 6). The small G-protein Rab10 has a role in regulating LPS induced trafficking of TLR4 from the Golgi (7). Also, we have reported that emp24 domain containing-protein 7 (TMED7) associates with TLR4 in the Golgi and affects TLR4 signalling activity (2).

N-linked glycosylations begin in the ER and are then further trimmed and elongated in the Golgi. TLRs have multiple glycosylations and a few of these sites influence the transport and secretion of members of the TLR-family. Association of TLR4 with the co-receptor MD2 in the early secretory pathway is essential for proper glycosylation at Asn526 and Asn575. Only the properly glycosylated TLR4 (130 kDa) is secreted onto the cell surface whereas hypoglycosylated TLR4 (110 kDa) is not detected on the cell surface in MD2 deficent cells (8–10). The ectodomain of human TLR2 has four predicted N-glycosylation sites, and one residue in particular is critical for secretion. Site-specific mutagenesis of Asn442 drastically reduces the secretion levels of TLR2, implying that this residue is an important factor for proper TLR2 biosynthesis and trafficking. Secretion of TLR2 is affected less by mutation of the other three glycosylation sites (11). Human TLR3 is also highly glycosylated and two residues have been associated with regulation of TLR3 endosomal expression. These two residues, N196 and N247, are potential targets for N-glycosylation and mutagenesis causes a large reduction in expression levels (12).

Transport of proteins between membrane bound compartments occurs via a general mechanism, which involves formation of vesicles on the donor compartment, vesicle fission, tethering and finally fusion with the acceptor compartment. There are a few known classes of vesicle involved in transport between membrane bound compartments. One example is Clathrin-coated vesicles which mediate protein transfer between the trans-Golgi compartment, the plasma membrane and endocytic pathways (13). Another group of coated vesicles are coat-protein complex (COP) - coated vesicles that mediate bidirectional membrane transport between the endoplasmic reticulum (ER) and the Golgi. The anterograde transport of proteins from the ER through COPII coated vesicles is balanced by retrograde transport from the Golgi in COPI coated vesicles in a process that recycles vesicle components and retrieves ER resident proteins. The COPI/II machinery is mechanistically similar but molecularly distinct (14, 15).

Nascent secretory proteins are translated and translocated to the ER where they are folded and subjected to post-translational modifications which can include N-glycosylations, disulphide bond formation and glycophosphatidylinositol (GPI) anchoring. Properly folded secretory soluble proteins and integral membrane proteins are exported from the ER. ER export is the first step in a transport chain that will direct them to their proper cellular localisation such as storage vesicles, lysosomes, the Golgi complex and the cell surface.

Cargo proteins are segregated from ER resident proteins and selectively sorted into ER-derived secretory vesicles by the cytoplasmic coat protein complex II (COPII) machinery. COPII vesicle budding occurs at specialized subdomains on the ER membrane, ER exit sites (ERES). ERES forms localized punctate structures on the ER membrane and are enriched in proteins of the COPII machinery (10, 16–18). The COPII coat consists of five subunits, Sar1p, Sec23p, Sec24p, Sec13p and Sec31p. Most mammalian COPII subunits have one or several paralogues with partially unique functions. The subunits are sequentially recruited in the process of COPII coat assembly and required for formation of transport vesicles that carry selected cargo proteins from the ER to the Golgi (19, 20). Upon vesicle fusion, the cargo proteins can proceed to transport towards the Golgi or be recycled back to the ER membrane in coat protein complex I (COPI) vesicles (21–26).

The COPII proteins play an important role in cargo sorting as they recognize and export a wide range of secretory proteins, both integral membrane proteins and soluble luminal proteins. There are several known transmembrane sorting receptors active in anterograde and retrograde ER to Golgi protein transport, as members of the p24 (TMED) family. Members of this protein family all have a similar membrane topology, a single spanning transmembrane domain, a N-terminal lumen domain and a cytosolic C-terminal tail with either COPI and/or COPII sorting motifs. Some members of the p24 family have been shown to interact and influence the trafficking of a variety of secretory cargo proteins. For example, silencing of TMED2 and TMED10 in yeast delays the trafficking of GPI-anchored protein Gas1p and TMED2 is required for Gasp1 incorporation into COPII vesicles (26). In mammalians, silencing of TMED10 impairs the trafficking of the GPI-anchored protein DAF and partial silencing of TMED7 affects TLR4 innate immune signalling (2, 27). The transmembrane region of TMED2 has been shown to interact with the head-group of C18 sphingomyelin lipid, suggesting that a protein-lipid interaction might regulate p24 proteins active and inactive state (28).

It has been proposed that p24 proteins form heterotetrametric complexes with members from each subfamily. In yeast, TMED2 and TMED10 or TMED9 respectively were found to be required for an efficient ER to Golgi transport of a subset of secretory proteins. In humans, it has been reported that formation of heterotetrametric complexes, consisting of one member of each subfamily is necessary for proper function and subcellular localisation. In addition, efficient ER-export and proper subcellular distribution of human TMED7 in yeast requires co-expression of TMED2, 9 and 10 (29, 30). However, other studies in yeast predict that p24 proteins predominately exist as monomers or dimers in a dynamic equilibrium. Such studies suggest that all p24 proteins are present as heterodimers but the monomer-dimer stoichiometry distribution in different organelles varies between members. The subcellular localisation appears to influence their oligomeric state and hence potentially their functional role in different compartments (31). These studies further emphasize that all p24 protein appears to have a distinct functional role and that their oligomeric properties are essential to their function and characteristics. Here, we showed that TMED7 and TMED2 play a role in transport of TLRs between ER and Golgi. Our results suggest that TMED7 cooperates with TMED2 to transport TLR2 and TLR4 and that N-glycosylation has an important role in this process

## RESULTS

### TMED2 and TMED7 interact in vivo

To determine whether TMED7 cooperates with TMED2 to regulate cargo sorting and transport between the ER and Golgi, Proximity Ligation Assay (PLA) experiments were performed in HEK293T cells and murine bone marrow-derived macrophages (BMDM). Both TMED2 and TMED7 are endogenously expressed in HEK293T cells and BMDM cells. However, the lack of high sensitivity antibodies targeting these native proteins prompted the use of exogenously expressed protein constructs with suitable C-terminal tags. HEK293T cells were transiently transfected with TMED2 and either full-length TMED7 or its ectodomain (CC-FLAG plasmid), subjected to PLA treatment and visualised using confocal microscopy. Our results (Figure 1and Figure S1A-D) suggest that full-length TMED2 and TMED7 are in close proximity resulting in PLA signals. The PLA signals are not localized within the nucleus but in other non-stained intracellular compartments. The interaction between TMED7 and TMED2 was confirmed by co-transfection with TMED7 ectodomain suggesting that the soluble form of TMED7 co-localises with the membrane attached form of TMED2, probably in the endoplasmic reticulum. These results suggest that TMED7 and TMED2 interact in HEK293T cells and that interaction between these two proteins only involves the TMED7 ectodomain. To validate the results, murine BMDM cells were transduced with lentivirus encoding TMED7 and TMED2, subjected to PLA treatment and visualised using confocal microscopy. Similar to results obtained in HEK293T cells, PLA signals were detected throughout the cells but not in the nucleus.

**Figure 1:**
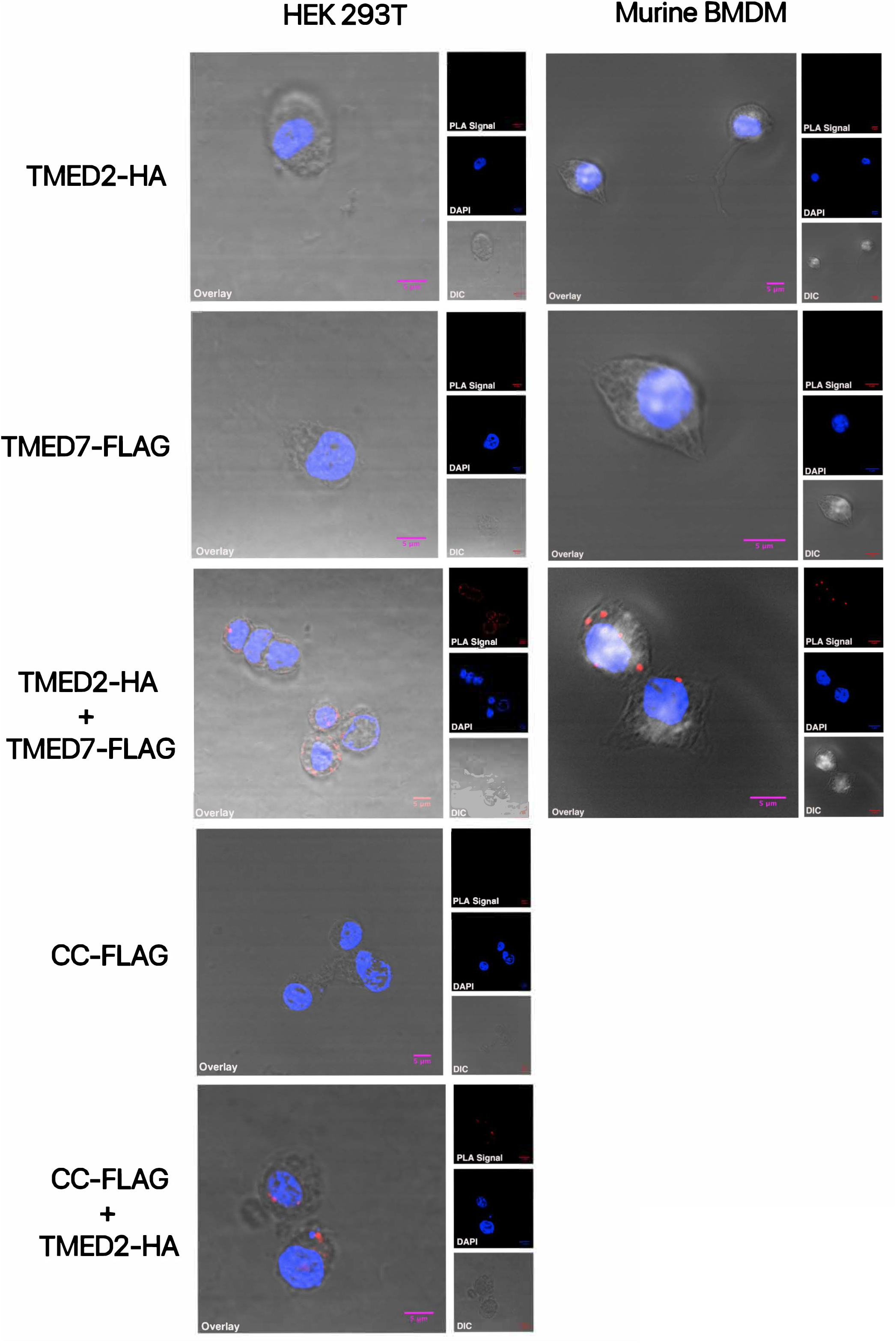
TMED2 interacts with TMED7. Proximity Ligand Assays (PLA) in HEK293T cells and murine BMDM enable visualisation of protein-protein interactions. Cells were transiently co-transfected and fixed prior to PLA staining and analysis by confocal microscopy. The fluorescent PLA signal was visualised in the red (559 nm) channel. The nucleus, stained with DAPI, was visualised in the blue (405 nm) channel, while images of cell structures were obtained using DIC microscopy. The overlay shows a merged, pseudo-coloured images of the three images collected with the above listed channels. Images are representatives of three independent experiments. PLA signals were produced in HEK293T cells and BMDM co-transfected with TMED2-HA and TMED7-FLAG, and HEK293T cells co-transfected with CC-FLAG (TMED7 ectodomain) and TMED2-HA. No PLA signal was detected in cells transfected with either TMED2-HA, TMED7-FLAG or CC-FLAG.

Together these results suggest that TMED7 and TMED2 interact and that the interaction does not require either the C-terminal tail nor the transmembrane region. These results are, partially, comparable to previous studies suggesting that the N-terminal GOLD domains of TMED2 and TMED5 interact weakly (32). However, the common hypothesis is that any homo- or hetero-oligomeric behaviour of p24 proteins involves the N-terminal coiled-coil domain. Further studies are required to identify the interaction interface and the molecular mechanism behind the formation of hetero-oligomeric p24 complexes. Also, the functional role of this interaction remains unknown.

### TMED2 and TMED7 Interacts with some TLRs

Our next step was to elucidate whether TMED2 and TMED7 cooperate to regulate cargo sorting and transport of Toll-like receptors by establishing if members of the TLR family interact with both cargo proteins. This was addressed by performing co-immunoprecipitation (co-IP) experiments to determine if TMED7 or TMED2 form complexes with members of the TLR family in HEK293T cells. Additionally, Proximity Ligation Assay (PLA) experiments were performed to visualise protein-protein interactions between TLRs and TMED7 or TMED2 in HEK293T cells.

First, HEK293T cells were co-transfected with either TMED7 or TMED2 along with one member of the TLR family. Forty-eight hours post-transfection the cells were lysed, forming the whole cell lysates used for co-immunoprecipitation, and analysed by Western blotting. In these assays both TLR2 and TLR4 but not TLR3 co-purified with TMED7. (Figure 2A). In contrast to TMED7, TMED2 co-immunoprecipitated with TLR2, TLR4 and TLR3 (Figure 2B). Together, these results indicate that TMED2 and TMED7 form complexes with some TLRs.

**Figure 2.**
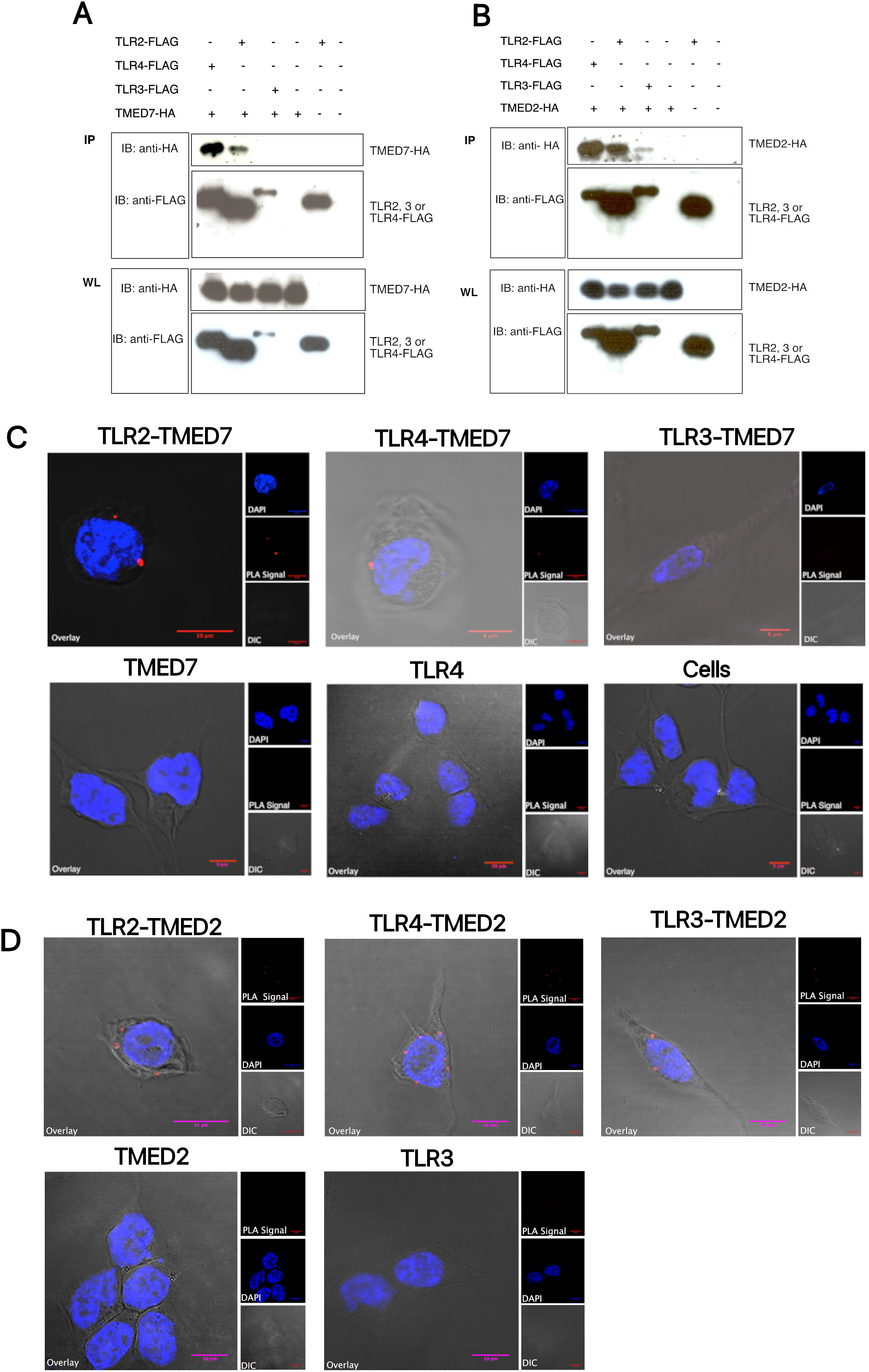
TMED7 Interacts with membrane TLRs and TMED2 with both endosomal and membrane. A and B: HEK293T cells were transiently transfected with the plasmids listed. Following cell lysis, the whole cell lysates were either prepared for analysis or incubated with M2 anti-FLAG beads to immunoprecipitate protein (IP) with FLAG-tags. Whole lysate (WL) and IP were resolved by SDS-PAGE and analysed by western blot using immunoblotting (IB). C and D: HEK293T cells were transiently co-transfected with plasmids listed and fixed prior to PLA staining and analysis by confocal microscopy. The PLA signal was visualised in the red (559 nm) channel. The nucleus stained with DAPI was visualised in the blue (405 nm) channel. Images of cell structures where obtain using DIC microscopy. The overlay shows a merged, pseudo-coloured images of the three images collected with the above listed channels. Images are representatives of three independent experiments. Scale bar 5 or 10 μm.

While these results suggest that TMED7 and TMED2 interact with a distinct subset of TLRs, they do not clarify whether the interaction is direct or indirect involving bridging interaction partners. To address this issue, the interaction between TLRs and p24 proteins was investigated using proximity ligation assays (PLA). HEK293T cells transiently co-transfected with TMED7 and either TLR2, TLR4, TLR5, TLR3 or TLR9 are shown in Figure 2C and Figure S1E-F. These results suggest that TMED7 interacts with TLR2, TLR4 and TLR5 which all belong to the subgroup of TLRs that predominately signal from the cell surface. In contrast, no PLA signals were produced in cells co-expressing TMED7 with either TLR3 or TLR9, receptors that signal from endosomal compartments in response to viral nucleic acids.

PLA was also used to investigate whether TMED2 interacts with members of the TLR family. The results in Figure 2D and Figure S1G-H, show that TMED2 interacts with TLR2, TLR4 and TLR3. These observations are supported by the results from co-IP, suggesting that TMED2 interacts with both endosomal TLR3 and with cell surface TLR2 and TLR4, while TMED7 is selective for TLR2, TLR4 and TLR5. We showed previously that TMED7 co-immunoprecipitates with TLR4 but not with TLR3 (2) which we have confirmed here. Interestingly, we also find that TMED7 is not exclusive for TLR4 but also interacts with TLR2 and TLR5. In contrast, TMED2 does not appear to be selective for the subgroup of TLRs signalling from the cell surface as it also interacts with endosomal TLR3. Taken together, these results provide evidence that TLRs and TMEDs interact but further work is required to elucidate whether TMED2 and 7 act as sorting adaptors for transport of TLRs between the ER and Golgi.

### TMED2 and TMED7 Affects the ER Export of some TLRs

As TLRs lack the C-terminal di-phenylalanine (FF) sorting motif required for a direct interaction with COPII subunits it is likely that adaptor proteins such as the TMED family are required for trafficking from the ER. Given that TMED7 and TMED2 interact with some TLRs, it is possible that these p24 proteins cooperate to regulate ER to Golgi transport of TLRs.

Most, though not all, members of the TMED family have the FF sorting motif which is responsible for interacting with COPII subunits. Removal of the FF motif disrupts the interaction with COPII subunits and results in TMED proteins and associated cargo being retained in the ER. To explore whether TMED7 and TMED2 are required for ER to Golgi transport of TLRs, HEK293T cells were transfected with either full-length or truncated p24 constructs. The cells were fixed and visualised using confocal microscopy. Prior to analysis by confocal microscopy, the ER and the Golgi were immunostained with primary anti-PDI antibody and an anti-Giantin antibody respectively, followed by secondary Alexa Fluor conjugated antibodies. Pearson’s correlation coefficient (R-values) for the whole image (IMAGE) and for the Region of Interest (ROI) were calculated. This method measures the intensity correlation, the degree of co-localisation, between each component of a dual colour image. R values ranges from −1 to +1, where the result is +1 for perfect correlation, 0 for no correlation and −1 for perfect anti-correlation.

In Figure 3A, HEK293T cells were transfected with either TMED7-mCherry (top) or TMED7T-mCherry (bottom). Overlay images are a merge of the red channel with either the far-red (ER) or with the blue (Golgi), R-values represent an estimate of the degree of correlation between the two respective dual-colour images. The merged images in the left top panel, indicate that full-length TMED7 does not co-localise well with the ER, which is supported by the R-values of −0.56 and −0.28. Images in the right top panel, indicate that full-length TMED7 is mostly localised in the Golgi exhibiting positive R-values of 0.45 and 0.88 respectively. The images in panel left bottom show that truncated TMED7T co-localised to a high degree with the ER (positive R-value of 0.80). In contrast, images in panel right bottom show that truncated TMED7T do not co-localise with the Golgi which is supported by the negative R value of −0.35. These results were confirmed using murine BMDM cells co-transduced with lentivirus encoding either TMED7-HA and TMED7T-HA (Supplementary Material Figure S2A). Together, these results suggest that removal of the C-terminal FF sorting motif retains TMED7 in the ER.

**Figure 3.**
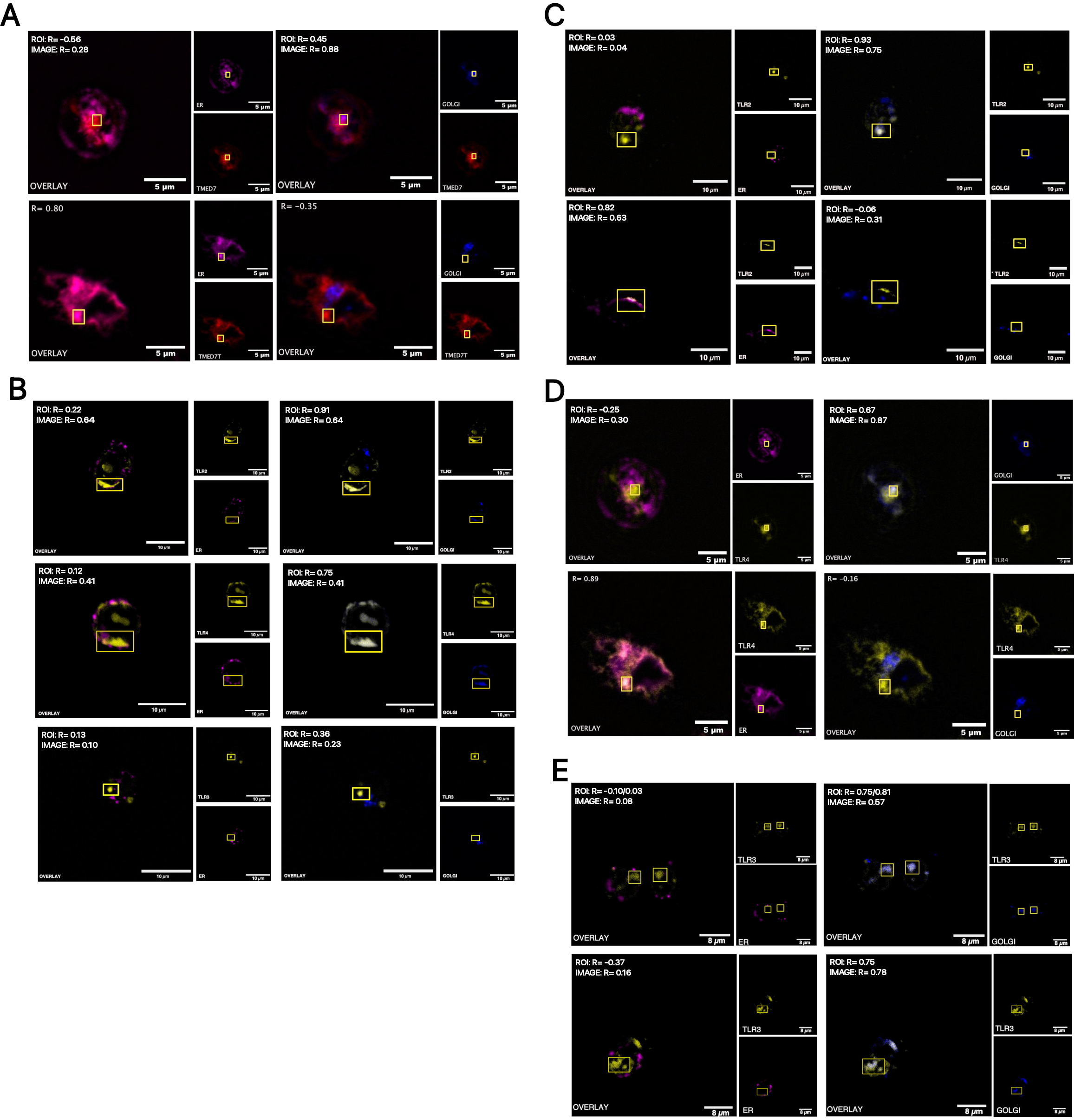
TMED7 Affects the ER export of some TLRs. Triple-colour imaging of HEK293T cells transiently expressing, A: TMED7-mCherry or TMED7T-mCherry, B: TLR2-EYFP, TLR4-Citrine and MD2, or TLR3-EYFP, C: TLR2-EYFP and TMED7-mCherry or TMED7T-mCherry, D: TLR4-citrine and TMED7-mCherry or TMED7T-mCherry, E: TLR3-EYFP and TMED7-mCherry or TMED7T-mCherry. TMED7 constructs were visualised in the red (559 nm) channel and TLR constructs in the yellow (515 nm) channel. The ER was immunostained and visualised in the far-red (635 nm) channel, while the Golgi in the blue (405 nm) channel. The overlay is a marge of the red channel with either the far-red or the blue channel respectively. Quantification of the Pearson’s coefficient (R) for the dual colour merged imaged. Images are representatives of three independent experiments. Panel A and D: scale bar 5 μm; panel E; scale bar 8 μm; panel B and C: scale bar 10 μm.

After confirming that removal of the FF motif retains the p24 cargo sorting receptors in the ER, the subcellular localisation of TLR2, TLR3 and TLR4 in HEK293T cells with an undisrupted ER-Golgi transport network was investigated. HEK293T cells were transiently transfected with constructs encoding either TLR2-EYFP, TLR3-EYFP or TLR4-Citrine respectively. The subcellular localisation of TLR2 in HEK293T cells are presented in the Figure 3B (top panels). The images in the right top panel indicate that TLR2 mostly localised in the Golgi (R values of 0.91 and 0.64). However, some TLR2 was localised in the ER (left top) and another fraction in other undefined subcellular localisations. In Figure 3B, middle panel, images of TLR4 expressed in HEK239T cells are shown and the degree of co-localisation with the ER (left) and the Golgi (right) respectively are examined. These images suggest that TLR4 is mostly localised in the Golgi, however TLR4 does also partially co-localise with the ER and some TLR4 is present in other non-labelled parts of the cell. In contrast to TLR2 and TLR4, images in bottom panel indicate that TLR3 does not co-localise with the ER (left) or the Golgi (right) but instead is predominantly distributed in other intracellular compartments distinct from the ER-Golgi membrane complex.

To establish the role of TMED7 in the ER to Golgi transport of TLRs, the subcellular localisation of TLRs in HEK293T cells co-transfected with either full-length or truncated TMED7 (TMED7T) were visualised using confocal microscopy. In Figure 3C, HEK293T cells were co-transfected with TLR2-EYFP along with either full-length (top) or truncated (bottom) TMED7. The results show that when TLR2 was co-transfected with full-length TMED7, TLR2 predominantly localised in the Golgi (top right) although some TLR2 was found in other unlabelled subcellular regions. On the other hand, TLR2 displayed a low degree of co-localisation with the ER (R= 0.03-0.04). When co-transfected with truncated TMED7, TLR2 mostly localised in the ER (bottom left), exhibiting a high degree of co-localisation (R=0.82 and 0.63). Meanwhile TLR2 did not localise in the Golgi (bottom right). The results show that when co-transfected with ER-export deficient truncated TMED7, the ER to Golgi transport of TLR2 is altered causing accumulation of TLR2 in the ER.

Images presented in Figure 3D show HEK293T cells co-transfected with TLR4-Citrine and either TMED7-mCherry (panel top) or TMED7T-mCherry (panel bottom). From these images it is evident that when co-transfected with full-length TMED7, TLR4 co-localises to a high degree with the Golgi (top right) and only a small fraction is localised in the ER (top left). However, when co-transfected with truncated TMED7 (bottom left) TLR4 mostly localised in the ER exhibiting high correlation (R= 0.89), while almost no TLR4 localised in the Golgi (bottom right). These results imply that in the presence of ER-export deficient truncated TMED7, TLR4 fails to translocate from the ER to the Golgi suggesting that TMED7 is required for ER export of TLR4.

The results in Figure 3E show the subcellular localisation of TLR3-EYFP when co-transfected with TMED7-mCherry (top) or TMED7T-mCherry (bottom) in HEK293T cells. The merged images indicate that TLR3 when co-transfected with full-length TMED7 is predominantly localised in the Golgi (top right) (R= 0.57-0.81) and that co-localisation with the ER is limited (top left). In contrast to TLR2 and TLR4, co-transfection with truncated TMED7 does not retain TLR3 in the ER (bottom left) but instead is mostly localised in the Golgi (bottom right) (R=0.75-0.78). These results imply that TMED7 is not required for ER to Golgi transport of TLR3, and this is supported by the observation that TMED7 failed to interact with TLR3. Given that TMED7 interacts with TLR2 and TLR4 but not with TLR3, these results further support the observation that TMED7 might regulate the ER to Golgi transport of TLR2 and TLR4 but not of TLR3. These results were validated using murine BMDM cells co-transduced with lentivirus encoding either TMED7-HA and TMED7T-HA along with TLR3-Citrine or TLR4-Citrine (Supplementary Material Fig. S1B).

After establishing the role of TMED7 in the ER to Golgi transport of some TLRs, the question regarding the role of TMED2 in the ER to Golgi transport of TLRs was investigated. In figure 4A, HEK293T cells were transiently transfected with either TMED2-mcherry (top) or TMED2T-mcherry (bottom). In panel top left, the images indicate that full-length TMED2 is predominantly localised in the ER and the positive R-values of 0.60 and 0.68 suggest that TMED2 colocalises to some extent but not fully with the ER. In panel top right, the images imply that the degree of co-localisation between TMED2 and the Golgi is low, with R-values of 0.10 and 0.34. Similarly, truncated TMED2T is mostly localised in the ER (bottom left) but was not present in the Golgi (bottom right). Interestingly, both full-length and truncated forms of TMED2 appear to predominantly localise in the ER making it difficult to assess whether removal of the FF motif resulted in a reduced level of TMED2 ER export. However, truncated TMED2 exhibited a higher degree of intensity correlation with the ER than full-length TMED2, with R-values of 0.74 and 0.80 versus 0.60 and 0.68 respectively, which might suggest that removal of the FF motif promotes accumulation of TMED2 in the ER. To confirm these observations, murine BMDM cells were co-transduced with lentivirus encoding either TMED2-HA and TMED2T-HA (Supplementary Material Fig. S1B). TMED2-HA does not localise in the ER, with a low positive intensity correlation (R-value of 0.07), but instead mostly in other undefined compartments. In contrast, truncated TMED2-HA is predominantely localised in the ER with positive R-values of 0.94, suggesting a high degree of intensity correlation between the observed channels.

**Figure 4.**
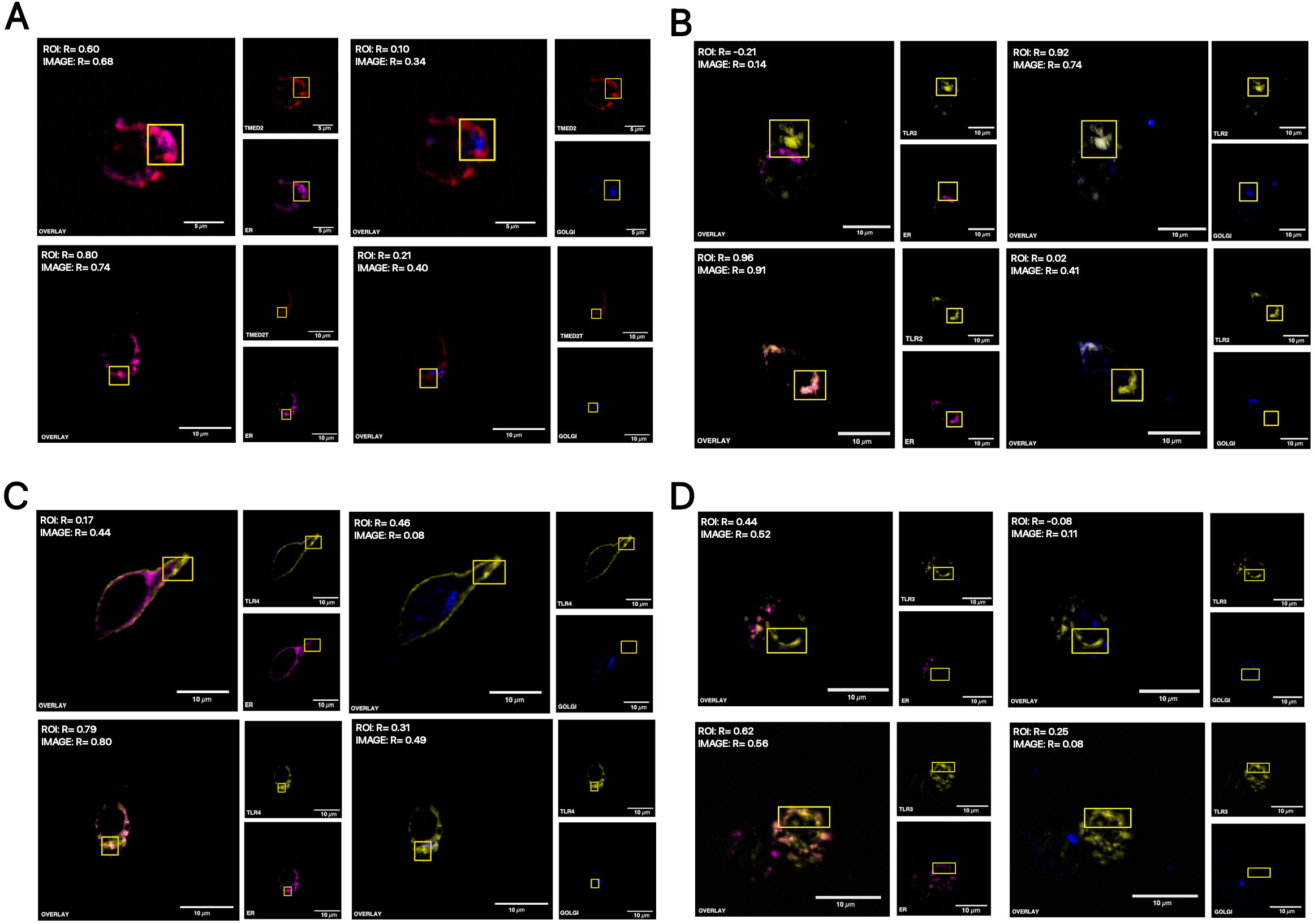
TMED2 regulates the ER Export of some TLRs. : Triple-colour imaging of HEK293T cells transiently expressing, A: TMED2-mCherry or TMED2T-mCherry; B: TLR2-EYFP and TMED2-mCherry or TMED2T-mCherry; C: TLR4-citrine and TMED2-mCherry or TMED2T-mCherry, D: TLR3-EYFP and TMED2-mCherry or TMED2T-mCherry. TMED2 constructs were visualised in the red (559 nm) channel and TLR constructs in the yellow (515 nm) channel. The ER was immunostained and visualised in the far-red (635 nm) channel, while the Golgi in the blue (405 nm) channel. The overlay is a marge of the red channel with either the far-red or the blue channel respectively. Quantification of the Pearson’s coefficient (R) for the dual colour merged imaged. Images are representatives of three independent experiments. Scale bar 10 μm.

To examine whether TMED2 is involved in the ER to Golgi transport of TLRs, confocal microscopy was used to assess the subcellular localisation of TLRs in HEK293T cells co-transfected with either full-length or truncated TMED2 constructs. Figure 4B shows HEK293T cells co-transfected with TLR2-EYFP and either TMED2-mCherry (top) or TMED2T-mCherry (bottom). In the top left panel, the merged image indicates that TLR2 does not localise in the ER when co-transfected with full-length TMED2. Instead, TLR2 mostly localised with the Golgi (top right), displaying a high degree of intensity correlation between the two merged channels (R-values of 0.94 and 0.74). The results suggest that over-expression of full-length TMED2 does not affect the ER to Golgi transport and subsequently the subcellular localisation of TLR2. Images in the bottom panel show that the presence of full-length TMED2 containing the FF-motif is relevant for ER exiting of TLR2. TLR2 when co-transfected with truncated TMED2 predominantly localised in the ER (bottom left), though a small fraction of TLR2 reached the Golgi (bottom right). Consequently, the results suggest that the presence of truncated TMED2 significantly reduces the level of TLR2 ER export.

The images in Figure 4C show that TLR4 when co-transfected with full-length TMED2 (top) predominantly localises in subcellular regions on or near the cell surface, while TLR4 co-localises poorly with both the ER and the Golgi. However, when co-transfected with truncated TMED2 (bottom), TLR4 mostly co-localises with the ER (R-values 0.79 and 0.80). However, a small but undefined fraction of TLR4 appeared to reach the Golgi (bottom right) showing that truncated TMED2 reduces the level of ER to Golgi transport of TLR4. This could be explained by an incomplete suppression of endogenous TMED2 facilitating transport of a small fractions of TLR4.

In Figure 4D, HEK293T cells were co-transfected with TLR3-EYFP along with either TMED2-mCherry (top) or TMED2T-mCherry (bottom). These results show that TLR3 when co-transfected with full-length TMED2 is primarily localised in non-labelled subcellular compartments and co-localise to a minor degree with the ER (top left) and the Golgi (top right). However, the subcellular localisation of TLR3 was different when co-transfected with truncated TMED2. In these cells, TLR3 exhibited a positive degree of co-localisation with the ER (bottom left) but not with the Golgi (bottom right), however an undefined fraction of TLR3 was also observed in non-labelled compartments. Thus, the results presented in these images imply that in the presence of truncated TMED2, TLR3 is partially but not completely retained in the ER. Suggesting that TMED2 might have role in the ER to Golgi transport of TLR3 but is not required for the ER export of TLR3. However, it could also be due to an incomplete suppression of endogenous TMED2. These results were also validated using murine BMDM cells co-transduced with lentivirus encoding either TMED2-HA and TMED2T-HA along with TLR3-Citrine or TLR4-Citrine (Supplementary Material Figure S2C).

### TLR2 N-Glycosylation State Affects the Level of ER Export

TMED7 and TMED2 interact with some TLRs, and presumably act as cargo sorting receptors regulating their transport from the ER to Golgi. However, it is still unclear how p24 proteins recognize mature cargo proteins and distinguish them from misfolded proteins or ER resident proteins. Given the wide range of secretory proteins that are processed and transported from the ER, it has been proposed that p24 proteins recognize common structural epitopes rather than unique features (33).

Attachment of glycans to asparagine (N) residues in the ER is a common and conserved process among eukaryotes (1, 34, 35). N-glycosylation occurs at asparagine residues linked to consensus sequence N-X-S/T, where X is any amino acid except proline (P). Members of the TLR family are all glycoproteins with multiple N-glycosylation sites. For example, human TLR2 has 4 N-glycosylation sites distributed in the N-terminal ectodomain. To determine whether the glycosylation status of TLR2 affects the level of ER to Golgi transport, the subcellular localizations of wild-type TLR2 and N-glycan mutants cells were analysed by confocal microscopy. HEK293T cells that do not express TLR2 were transiently transfected with plasmids encoding TLR2 wild type or different TLR2 mutants carrying one or two N-glycan sites with N-terminal FLAG tags (see (11)). To enable visualization by confocal microscopy, the cells were fixed and subjected to immunofluorescence staining. TLR2 WT was predominantly localized in intracellular regions excluding the ER and only a small amount of TLR2 appeared to co-localise with the ER (Figure 5A). Although, these results are from unstimulated cells, some wild-type TLR2 localized in regions near or on the cell surface.

**Figure 5.**
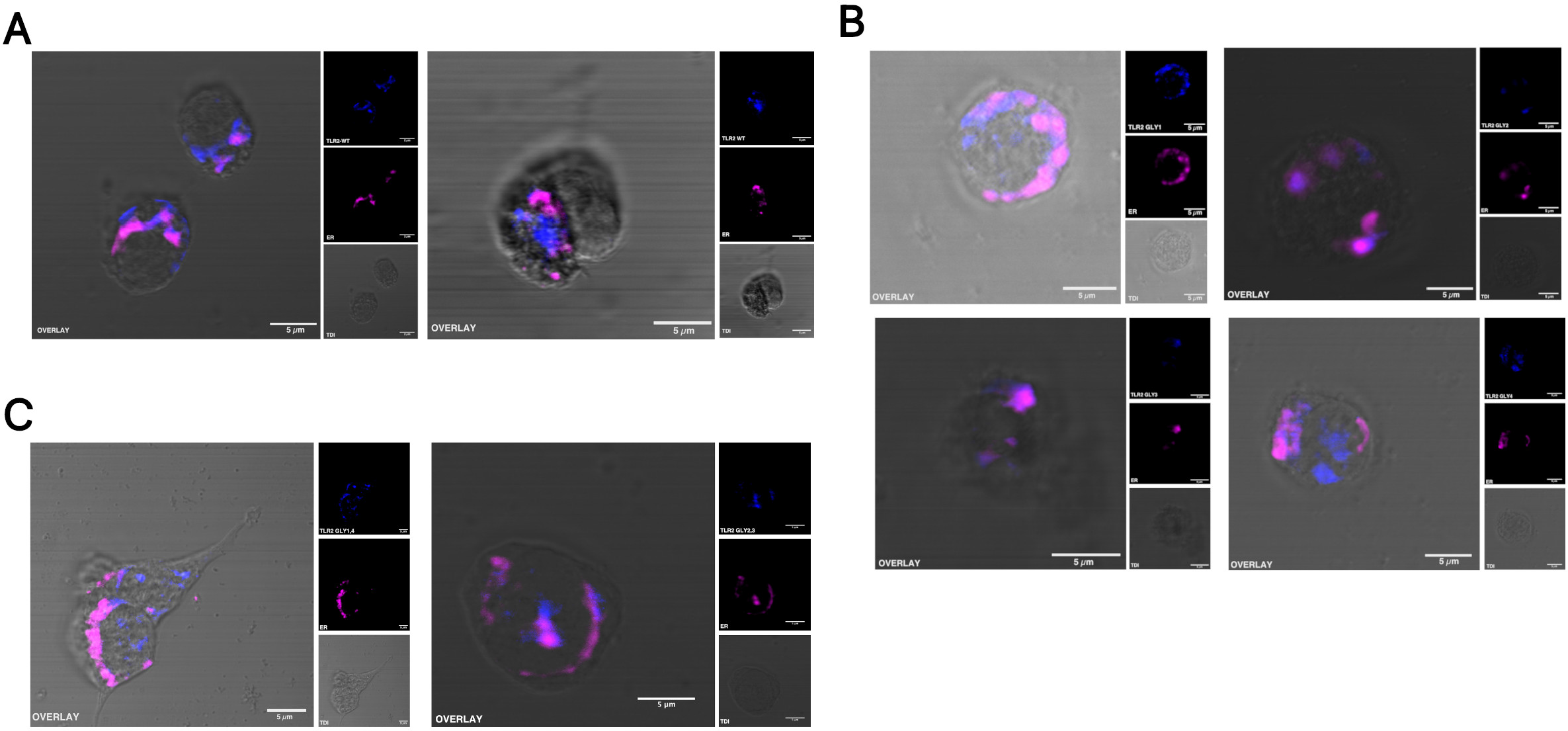
TLR2 N-Glycosylation State is necessary for ER export. The subcellular localisation of TLR2 WT (A) and TLR2 mutants expressing one glycosylation (B) or two glycosylations (C) in HEK293T cells, was visualized using confocal microscopy. TLR2- and TLR2GLYm-FLAG, immunostained with anti-FLAG antibody was visualized in the far-red (635 nm) channel, while the ER, immunostained with anti-PDI antibody was visualized in the blue (405 nm) channel. Images of cell structures were obtain using DIC microscopy. Overlay shows a merged, pseudo-colored images of the three images collected with the above listed channels. Mounting medium was added prior to analysis by confocal microscopy. Images are representatives of three independent experiments. Scale bar 5 μm.

In Figure 5B, HEK293T cells were transiently transfected with plasmids encoding TLR2 containing only one N-glycan site, either N114(GLY1), N199(GLY2), N414(GLY3) or N442(GLY4). In panel top left, HEK293T cells were transiently transfected with TLR2GLY1, subjected to immunofluorescence staining and then analysed by confocal microscopy. The results show that TLR2GLY1 mostly co-localized with the ER, but the protein was also detected, to a lesser extent, in other intracellular regions. In contrast, TLR2GLY2 (top right) and TLR2GLY3 (bottom left) were both predominantly localized in the ER and barely detected in other areas of the cells. Another TLR2 glycan mutant, TLR2GLY4, was transiently transfected in HEK293T cells and visualized by confocal microscopy (bottom right). Here, TLR2GLY4 co-localized poorly with the ER but was instead abundant in other subcellular localizations. All together, these results imply that glycosylation in N442 is important for ER export as the level of TLR2 ER export is significantly reduced for receptors missing this N-glycan.

Next, HEK293T cells were transiently transfected with TLR2 plasmids containing two of the four N-glycan sites, stained for immunofluorescence and analysed by confocal microscopy. The results are presented in Figure 5C, showing that TLR2GLY2,3 was predominantly localized in the ER (right) and in contrast to TLR2GLY1,4 that was mostly localized in subcellular regions distinct from the ER (left). TLR2 carrying glycan in N199 and/or N414 were mostly abundant in the ER implying that these TLR2 mutants have a reduced ER anterograde transport. In addition, TLR2 mutants expressing glycans in N442 (TLR2GLY4) and N442 plus N114 (TLR2GLY1,4) both exhibited a subcellular distribution similar to TLR2 wild-type (WT), which suggests that N-glycan in position N442 might play an important role in the transport of TLR2 from the ER. Interestingly, these findings are supported by previous studies showing that TLR2 cell surface secretion is reduced for TLR2 mutants lacking GLY4 (11).

Although, these results suggest that the N-glycosylation state of TLR2 influences the level of ER export, the in-cell situation may be more complex. These results are from unstimulated cells and the TLR2 distribution might alter upon receptor stimulation. Additionally, TLR2 is not endogenously expressed in HEK293T cells and consequently, these results might not represent native TLR2 expression. However, under these circumstances the subcellular distribution of various TLR2 N-glycan mutants were notably different compared to what was observed for wild-type TLR2, and it is evident that the level of TLR2 ER export in HEK293T cells is affected by N-glycosylation state. In conclusion, the data presented suggests that TLR2 N-glycosylation in position N442 is important for the transport of TLR2 from the ER.

## DISCUSSION

Earlier studies to investigate the functional role of p24 proteins have mostly been in yeast cells whose secretory pathway is less complex than in higher organisms and consequently their precise functional roles in mammalians is largely unknown. The available evidence suggests that p24 proteins function as cargo sorting receptors that regulate sorting and transport of cargo glycoproteins between the ER and the Golgi. However to date only a few cargo proteins have been identified and among them are members of the family of GPI-AP. Members of this family of proteins are attached to the membrane by a C-terminal glycolipid (GPI). Recent studies have showed that TMED2 recognises mature GPI-AP in the ER lumen and facilitates incorporation into COPII vesicles (36, 37). Also, previous work has shown that TMED7 is required for trafficking of TLR4 (2).

Altogether, the results from both immunoprecipitation assays and PLA suggest that TMED7 selectively associates with cell surface signalling receptors TLR2 and TLR4, while TMED2 in addition to TLR2 and TLR4 also interacts with TLR3 signalling from endosomal compartments. These results were obtained from cells overexpressing the various target proteins and it is possible that the protein-protein interactions found might not occur under steady state conditions in the cells. However, the results from two independent methods were consistent with each other and this enhances the reliability of our findings.

In our previous study (2) TMED7 knockdown enhanced the TRIF-dependent response of TLR4 after LPS stimulation without affecting the NF-κB response. These results were partially confirmed by studies in THP-1 cells transfected with siRNA for TMED7 which found a reduced amount of interleukin-6 (IL-6) and tumor necrosis factor-α (TNF-α), both activated by NF-κB, without affecting RANTES, an interferon-sensitive response element (ISRE)-dependent gene activated by the TRIF pathway. So, taken together these results suggest that TMED7 not only participates in the transport of cell surface TLRs but also supports their expression at the membrane. In this regard a patient has been identified with a homzygous deletion of TMED7 (Arg140*). The affected individual has a complex phenotype including Stevens-Johnson syndrome, a severe inflammation of the mucous membranes, that might be caused by activation of mislocalised TLRs (38). This pleiotropic phenotype also includes early onset visual impairment and is consistent with TMED7 functioning in vesicle trafficking of multiple proteins.

Although more studies are required to understand the precise mechanism by which p24 proteins recognise mature cargo proteins, our results show that TMED7 and TMED2 cooperate with an overlapping subset of TLRs to regulate anterograde trafficking. Our results are also consistent with previous studies showing that hetero-oligomers of TMED2 with other p24 family proteins are required to transport TLR4 and TLR2 and that association of TMED2 with other TMED proteins regulate their stability, localization and transport (31, 39–41). Experiments using mammalian and yeast cells already showed that TMED2 forms complexes with TMED10, TMED7 and TMED9, and these complexes are important for stability (29, 42).

To further explore the possible role of TMED7 and TMED2 in ER to Golgi transport of TLRs, we used dominant negative truncations of TMED 2 and 7 lacking the FF sorting motif and assayed for co-localisation in of TLRs (42). These experiments confirm that the FF motif is required for ER export and anterograde transport to the Golgi and that when co-expressed with ER export deficient TMED7, TLR2 and TLR4 were predominantly localised in the ER while TLR3 was trafficked to the Golgi. By contrast in the presence of full-length TMED7, TLR2, TLR3 and TLR4 were efficiently exported from the ER. Additionally, dominant negativeTMED2 significantly reduced the ER to Golgi export of TLR2 and TLR4, but also of TLR3. Interestingly dominant negative TMED2 significantly reduced but did not block ER export of TLR2, TLR3 and TLR4. The incomplete block of TLR transport in TMED2 ER export deficient cells could be explained in several ways. For example, there could be another protein with a function which overlaps that of TMED2 with the ability to partially restore the transport, but it could also reflect an incomplete suppression of native TMED2.

Together, the results suggest that TMED7 and TMED2 associates with partially overlapping subsets of TLRs and facilitate their incorporation into COPII vesicles and subsequent ER export. Although, there is a general consensus that p24 proteins act as cargo sorting receptors in the early secretory pathway at this point it is still unclear how p24 proteins recognise mature cargo proteins, such as TLRs, and incorporate them into COP transport vesicles for ER anterograde transport. However, due to the wide range of cargo proteins that are processed and transported throughout the early secretory pathway, it has been suggested they recognise structural epitopes, such as N-glycosylation, rather than unique features. For example, it was reported that TMED2 recognizes mature N-glycan of GPI-anchored proteins and that improperly glycosylated GPI-anchored proteins failed to be transported from the ER (33). Here the effect of the N-glycosylation state on the level of ER export of TLR2 was studied. Of the four N-glycosylation sites in TLR2 only N442 plays an important role for recognition and transport from the ER. However, although TLR2 N442S (GLY4) mutants mostly were retained in the ER, small traces of the mutant receptor were detected in other subcellular localisations. This could be because these TLR2 mutants are still recognised by the cargo sorting receptor but at a lower affinity than TLR2 carrying GLY4.

These results and previous work (11) suggest that the TLR2 N-glycosylation state influences the level of TLR2 ER export. Given that the N-glycosylation state affects the ER export of TLR2, it would be of interest to investigate whether it also affects the level of ER export for other TLRs. In addition, the results reveal a functional link between the N-glycan state and ER export of TLR2, where the presence of particular N-glycans appears to be required for an efficient ER export. This suggests that N-glycan maturation acts as a quality control system to prevent premature ER export. However, although it is clear that the N-glycan state of TLR2 is important for ER export, it remains to be addressed whether the N-glycan state of TLR2, and possibly of other TLRs, affects their association with TMED7 or TMED2. Another unresolved issue is the molecular mechanisms by which p24 proteins associate and disassociate with cargo glycoproteins such as TLRs. It has been suggested that cargo remains bound to the cargo sorting receptor until reaching ERGIC or the Golgi. For example, the cargo sorting receptor Erp44 associates with cargo proteins in the Golgi in a pH dependent manner to retrieve them back to the ER (43). A similar observation has been made for another cargo sorting receptor ERGIC-53, which associates with selected cargo at neutral pH in the ER whereas the more acidic environment in the ERGIC promotes dissociation (44).

Trafficking regulated by TMED2 is also important for innate immune responses to nucleic acid. Cytosolic double stranded DNA released by infection with herpes simplex virus generates the cyclic dinucleotide cGMP-AMP (cGAMP) by activating cGAMP synthase. cGAMP then binds to the intrinsic membrane protein stimulator of interferon genes (STING) which then interacts with TMED2 and is trafficked to the Golgi. Golgi resident STING/TMED2 complexes then recruit Tank-binding kinase 1 (TBK1) and IκB kinase-β that in turn activate NFκB and interferon response factor 3 (IRF3) (45). In conclusion specific anterograde transport mediated by the TMED family plays a central role in regulating innate immunity.

## MATERIALS AND METHODS

### Mammalian cells

HEK293T cells were cultured in complete medium (Dulbecco’s modified Eagle medium (Gibco) supplemented with Penicillin (100U/mL) and Streptomycin (100 μg/mL) (Gibco), and 10 % (v/v) heat-inactivated Fetal Bovine Serum (Sigma-Aldrich)). For lentiviral production, complete DMEM supplemented with 10 mM HEPES was used. The cell cultures were maintained at 37°C and 5% CO2 in a static incubator. The cells were grown in 75cm^2^ tissue culture flasks until they reached 60-70% confluency before being passaged and cells with passage numbers 2 to 15 were used for experiments. The cells were passaged by aspirating old medium, washed with PBS (Sigma), and detached by adding TrypLE Express (Gibco) following incubation for 2 min at room temperature. The cells were then resuspended using an equal volume of complete medium, pelleted at 250 g for 5 min and then resuspended in complete medium. Trypan Blue, an automatic cell counter (Nexelom Bioscience) and Cellometer cell counting chambers (Nexcelom Bioscience) were used to measure cell counts and cell viability. Cells were seeded at concentrations between 100000-150000 cells/μL.

Murine BMDM (Bone Marrow Derived Macrophages) cells were a gift from Dr Lee Hopkins, Department of Veterinary Medicine, University of Cambridge. The cells were cultured in complete DMEM supplemented with 10 mM HEPES (ThermoFisher) as described above. Cells were seeded at concentrations between 130000-150000 cells/mL and maintained at 37°C and 5% CO2 in a static incubator. The BMDM cells were passaged 2-5 times before being infected with lentiviral particles.

### Microorganisms

*E. coli* strains DH5α (Invitrogen) or DH10β (NEB) chemically competent cells were used for cloning experiments to generate desired constructs as well as for general plasmids transformations. Plasmids were subjected to antibiotic selection using Ampicillin or Carbenicillin at 100 μg/mL. For bacterial plasmid production, Luria-Bertani (LB) medium (10 gram/L Bactotryptone, 5 gram/L bacto-yeast extract and 10 gram/L NaCl in water) was used, or supplemented with 20 gram/L of agar for agar plates.

For lentiviral particles production, HEK293T cells were seeded on 12-well plates at 150000 cells/ml in complete DMEM supplemented with 10 mM HEPES. Before transfection the old medium was gently aspirated and replaced with new fresh complete DMEM supplemented with 10 mM HEPES (ThermoFisher). When reaching 70-80% confluency the cells were transfected with the corresponding pHR plasmid plus p8.91 and pMD-G plasmids (please, see recombinant DNA list in key resources table). Twenty-four hours post-transfection the medium was replaced with antibiotic free medium, then the cells were incubated for 48 hours when the medium containing lentiviral particles were harvested, centrifuged at 300g and filtered through a 0.22 μm filter. Lentiviral suspension was either used directly on target host cells or stored at 4°C for maximum one week before experiments.

### Co-immunoprecipitation

Co-immunoprecipitation (co-IP) was used to study protein-protein interactions in HEK293T cells. Plasmids encoding proteins of interest were transiently transfected into HEK293T cells in a 6-well format. Forty-eight hours post transfection cells were removed and incubated in 250 μL/well of HEK Lysis Buffer containing protease Inhibitor Cocktail Set V (Calbiochem) before being centrifuged at 13 000 rpm for 10 min. To immunoprecipitate proteins, 200 μL of supernatant was incubated with 20 μL of M2 anti-FLAG beads overnight at 4°C. Prior to incubation with proteins, the beads were equilibrated in TBS according to the manufacturer’s protocol. Following incubation, the anti-FLAG beads were washed 3 times with TBS to remove unbound proteins. Proteins were eluted from the beads by adding 4× LDS sample buffer and 1 mM DTT and boiled for 5 min. Prior to being loaded onto SDS-PAGE gels, protein samples were centrifuged at 1000g, 1 minute. Proteins were transferred to nitrocellulose membranes for protein detection by Western blot.

### Western blots

Proteins were transferred from SDS-PAGE gels onto nitrocellulose membranes (Amersham) in 1× standard transfer buffer at 15V for 1 hour. Membranes were blocked in 3% milk powder or 3% BSA in PBST for 1 hour at room temperature or overnight at 4°C. Membranes were probed with primary antibodies in PBST, washed 3 times for 5 min in PBST, and then probed with secondary antibodies. Membranes were washed 3 times for 2 to 5 min in PBS and incubated with Supersignal West Pro Plus Chemiluminescent Substrate for 5 min. For protein detection, films were exposed on Fuji Medical X-Ray Films (Fuji) in exposure times ranged from 10 s to 20 min. Films were developed using a Film Processor (AFP Imaging). Film exposure and developing were conducted in the dark.

### Confocal microscopy

HEK293T cells were seeded onto μ-Slide 8 well (Ibidi), upon reaching 30-40% confluency the cells were transfected with own shDNA amounts using PEI 25K transfection reagent. About forty-eight hours post transfection the medium was carefully aspirated, and cells washed with TPBS. Cells were fixed for 20 min at 4°C with 4% Paraformaldehyde and then permeabilized by incubation with 0.05% (v/v) Triton-x100 for 10 min. Cells were washed 3 times for 5 min with cold TBS. Cells blocked for 1 hour at room temperature in blocking buffer. Cells were then stained for immunofluorescence with primary antibodies (diluted 1:1000) towards Protein Disulfide Isomerase, PDI (ER-marker) and Giantin (Golgi-marker). Cells were then washed 3 times for 5 min with TBST and then incubated with Alexa Fluor conjugated secondary antibodies (diluted 1:1000 in antibody dilution).

For co-localisation analysis using BMDM cells, cells were induced with lentivirus encoding genes of interest and 4-5 days post lentiviral infection the cells were gently washed and aspirated, and then transferred to onto μ-Slide 8 wells (Ibidi). Cells were fixed by 20 min at 4°C with 4% paraformaldehyde and then permeabilized by 2 min incubation with acetone at −20°C. Cells were washed 3 times for 5 min with cold PBS. Cells blocked for 1 hour at room temperature in blocking buffer. Cells were then stained for immunofluorescence with primary antibodies (diluted 1:1000 in antibody dilution) towards PDI (ER-marker) and HA-tag (TMEDs-HA marker). Cells were then washed 3 times for 5 min with PBST and then incubated with Alexa Fluor conjugated secondary antibodies (diluted 1:1000 in antibody dilution). Cells were then washed 3 times for 2 min with PBST and allowed to dry in the dark. Samples were mounted for imaging by adding ProLong Diamond Antifade Mountant (Invitrogen) and allowed to cure overnight in the dark at 4°C before being visualized.

All microscopic experiments were conducted using an Olympus FV1200 Laser Scanning confocal microscope. LD lasers at 405 nm, 559 nm and 635 nm, and Argon Laser at 515 nmwere used to excite fluorochromes. The emission band settings were adjusted to capture maximal emission signal from a particular fluorophore while avoiding emission overlap from other fluorophores. Cells were visualized using a 60x oil immersion objective lens and a frame averaging of 5 was applied during image acquisition.

### Proximity Ligation Assay

HEK293T cells were seeded onto μ-Slide 8 well (Ibidi) and upon reaching 30-40% confluency the cells were transiently transfected with mammalian expression plasmids carrying the genes of interest. About forty-eight hours post transfection the medium was carefully aspirated and cells washed with PBS. Cells were fixed for 20 min at 4°C with 4% paraformaldehyde and then permeabilized by 2 min incubation with acetone at −20°C. Cells were washed 3 times with TPBS before being prepared for confocal microscopic analysis using Duolink^®^ Proximity Ligand Assay (PLA) in situ experiment.

In HEK293T cell, the Duolink^®^ In Situ Red Starter Kit Mouse/Rabbit (Sigma) was used according to manufacturer’s recommended protocol. Primary antibodies used were mouse anti-HA (Abcam) and rabbit anti-FLAG (Abcam). In murine BMDM cells, Duolink^®^ In Situ Red Starter Kit Goat/Rabbit (Sigma) was used according to manufacturer’s recommended protocol. On BMDM cells, the primary antibodies used were goat anti-DDDDK (Abcam) and rabbit anti-HA antibodies (Abcam). The samples were mounted for imaging by adding ProLong Diamond Antifade Mountant with DAPI (Invitrogen) and allowed to cure overnight in the dark at 4°C before being visualized.

### Differential Interference Contrast (DIC)

Along with each colored image visualized by confocal microscopy, in parallel an image of the cell structure was illuminated and collected using Differential Interference Contrast (DIC) Microscopy. This was enabled by selecting the DIC mode of the FV1200 Olympus Microscope. To illuminate the sample for DIC, the 635 nm or 488 nm laser was used at a low power.

### Quantification and statistical analysis

Images obtained from Confocal Microscopy were analysed and processed using Fiji ImageJ (https://fiji.sc) and co-localisation analysis was performed using plug-in coloc2. Images set for co-localisation analysis were subjected to rolling ball background subtraction of 1 prior to co-localisation analysis by coloc2. Calculation of co-localisation values were performed using Pearson’s coefficient (R). This method measures the intensity correlation, the degree of co-localisation, between each component of a dual color image. R values ranges from −1 to +1, where the result is +1 for perfect correlation, 0 for no correlation and −1 for perfect anti-correlation. Co-localisation analysis was performed either on the whole image and/or on a selected region of interest (ROI). Pearson’s coefficient (r) can range from −1 to 1 where a positive value indicates co-localisation (Bolte and Cordelieres, 2006). Pearson’s Correlation Coefficients are described under results sections when relevant and final figures were prepared using Affinity Designer and GIMP. PLA images were analysed using ImageJ plug-ins CellCounter and SpotCounter, A number (5 to 10) of representative images for each experimental condition were analysed and output data is presented in histograms that include the standard deviation (SD). Results from quantification PLA signals and mean PLA signal intensity are presented in supplementary material.

## AUTHOR CONTRIBUTIONS

J.E.J.H, S.G.S. and M.C.M. designed experiments. J.E.J.H. and M.F.S. performed microscopy analysis. J.E.J.H, S.G.S. and M.C.M analysed the data and wrote the manuscript with N.J.G.

## ACKNOWLEDGMENTS

We thank Dr. Lee Hopkins for helping on lentiviral transfection and BMDM cell culture.

---

**Figure S1:**
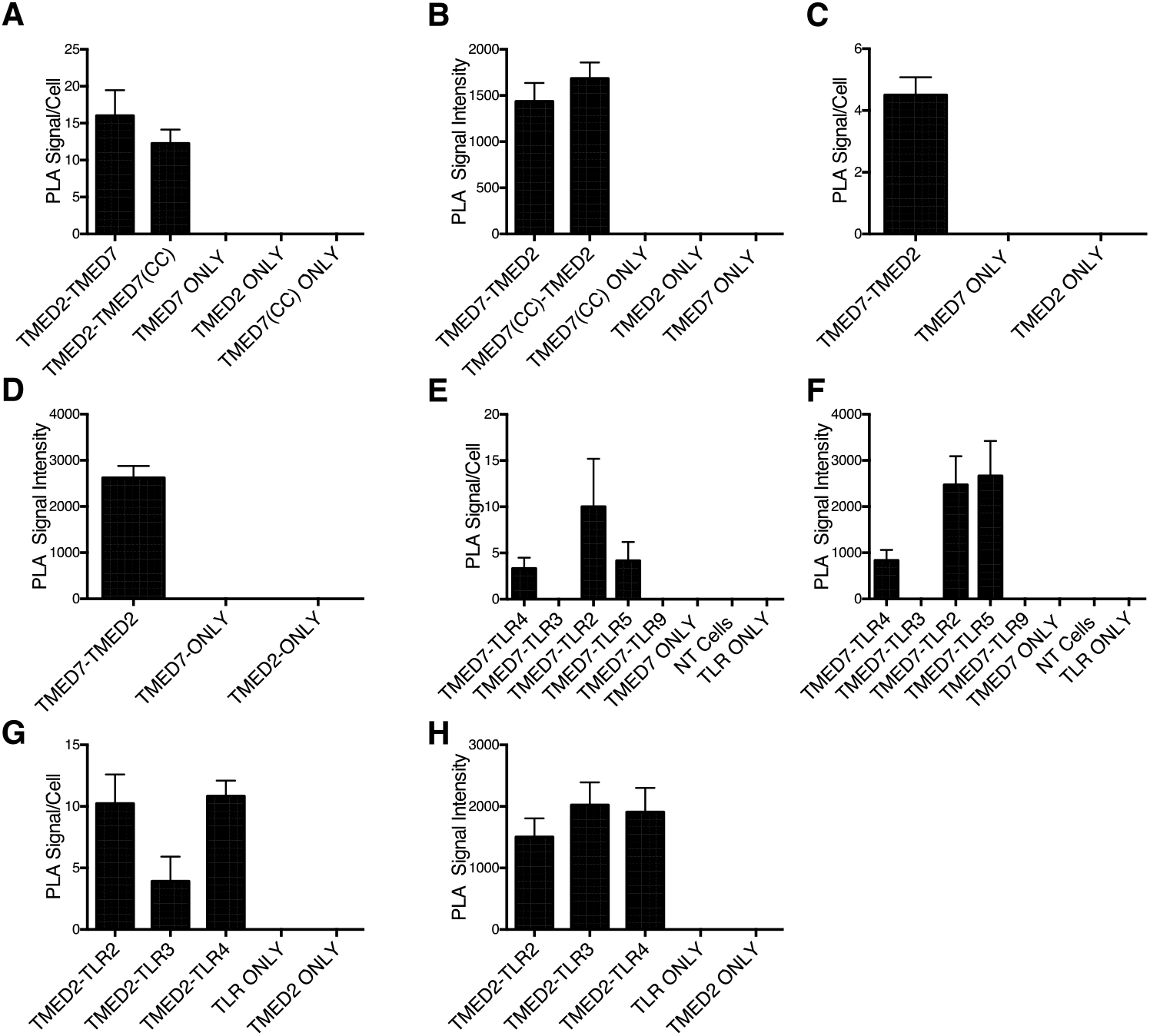
Quantification of PLA Signals. The quantification of the number of PLA Signal per cell (n) and mean PLA Signal Intensity (i), Relative Fluorescent Intensity, on the Y-axis is presented with the mean S.D. Cells were transfected or transduced with plasmids/virus encoding for constructs indicated on X-axis. A/B: TMED2 and TMED7 interact in HEK293T Cells. C/D: TMED2 and TMED7 interact in iBMBM cells. E/F: TMED7 interact with TLR2 and TLR4 but not with TLR3 in HEK293T cells. G/H: TMED2 interacts with TLR2, TLR4 and TLR3 in HEK293T cells. NT (non transfected cells), TLR only (cells transfected with either TLR2, TLR3 or TLR4).

**Figure S2:**
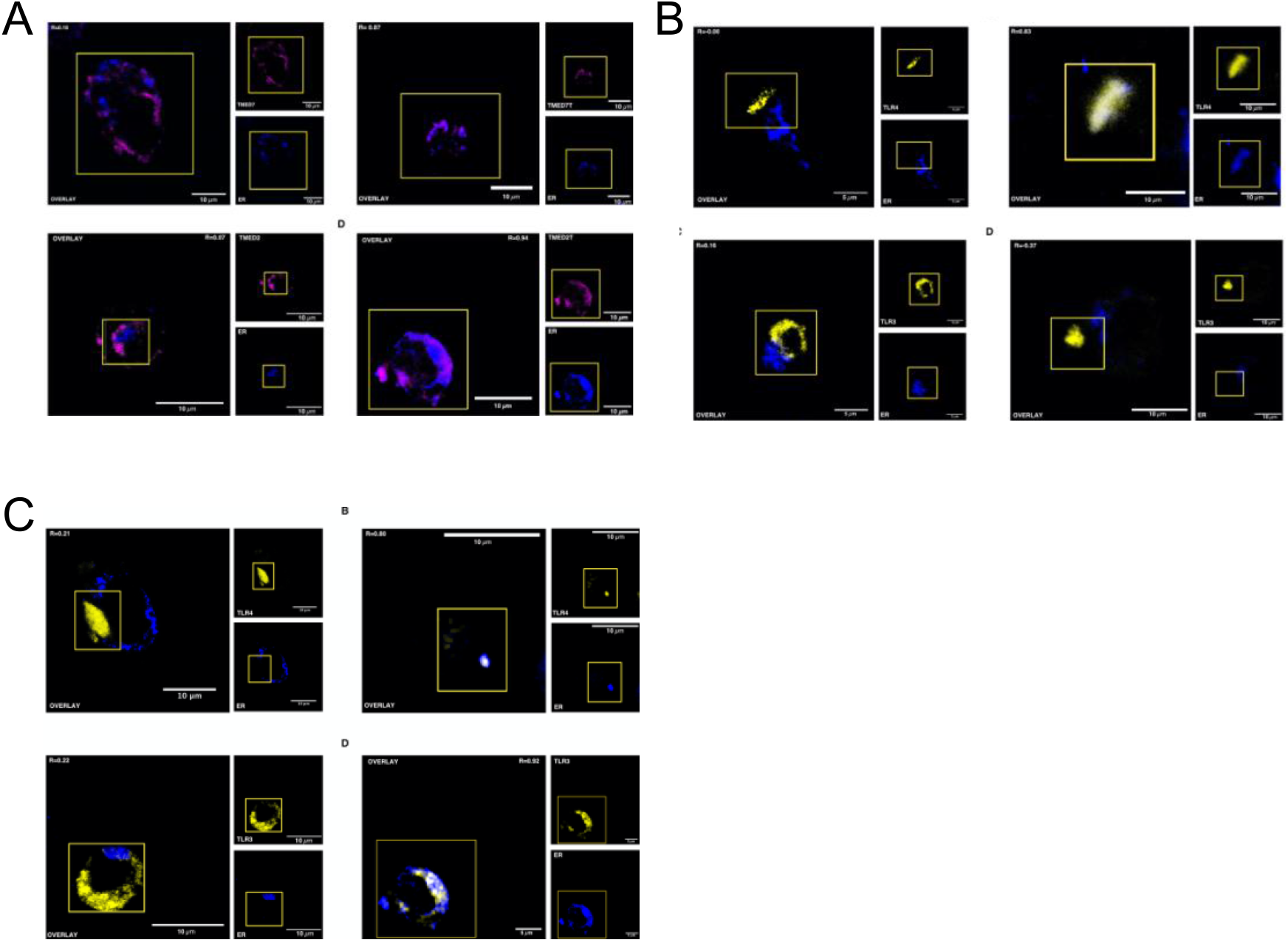
Removal of FF-motif retains both TMED2 and TMED7 in the ER and affects the ER Export of some TLRs in murine BMDM cells. Dual-colour imaging of murine BMDM cells transduced with lentivirus encoding, **A**: TMED7- and TMED7T-HA (top), and TMED2- and TMED2T-HA (bottom); **B**: TMED7-HA (top left) and TMED7T-HA (top right) with TLR4-citrine; TMED7-HA (bottom left) and TMED7T-HA (bottom right) with TLR3-citrine; **C**: TMED2-HA (top left) and TMED2T-HA (top right) with TLR4-citrine; TMED2-HA (bottom left) and TMED2T-HA (bottom right) with TLR3-citrine. HA-tag constructs were immunostained and visualised in the far-red (635 nm) channel and TLRs in the yellow (515 nm) channel, while the ER was immunostained and visualised in the blue (405 nm) channel. The overlay is a merge of the red channel with either the far-red or the blue channel respectively. Quantification of the Pearson’s coefficient (R) for the dual colour merged imaged. Images are representatives of three independent experiments. Scale bar 5 to 10 μm.

## REFERENCES

1. Lederkremer GZ. Glycoprotein folding, quality control and ER-associated degradation. Current opinion in structural biology. 2009;19(5):515–23.

2. Liaunardy-Jopeace A, Bryant CE, Gay NJ. The COP II adaptor protein TMED7 is required to initiate and mediate the delivery of TLR4 to the plasma membrane. Science signaling. 2014;7(336):ra70.

3. Lee BL, Moon JE, Shu JH, Yuan L, Newman ZR, Schekman R, et al. UNC93B1 mediates differential trafficking of endosomal TLRs. eLife. 2013;2:e00291.

4. Randow F, Seed B. Endoplasmic reticulum chaperone gp96 is required for innate immunity but not cell viability. Nature cell biology. 2001;3(10):891–6.

5. Takahashi K, Shibata T, Akashi-Takamura S, Kiyokawa T, Wakabayashi Y, Tanimura N, et al. A protein associated with Toll-like receptor (TLR) 4 (PRAT4A) is required for TLR-dependent immune responses. The Journal of experimental medicine. 2007;204(12):2963–76.

6. Wakabayashi Y, Kobayashi M, Akashi-Takamura S, Tanimura N, Konno K, Takahashi K, et al. A protein associated with toll-like receptor 4 (PRAT4A) regulates cell surface expression of TLR4. Journal of immunology. 2006;177(3):1772–9.

7. Wang D, Lou J, Ouyang C, Chen W, Liu Y, Liu X, et al. Ras-related protein Rab10 facilitates TLR4 signaling by promoting replenishment of TLR4 onto the plasma membrane. Proceedings of the National Academy of Sciences of the United States of America. 2010;107(31):13806–11.

8. Nagai Y, Akashi S, Nagafuku M, Ogata M, Iwakura Y, Akira S, et al. Essential role of MD-2 in LPS responsiveness and TLR4 distribution. Nat Immunol. 2002;3(7):667–72.

9. Wang Y, Chen T, Han C, He D, Liu H, An H, et al. Lysosome-associated small Rab GTPase Rab7b negatively regulates TLR4 signaling in macrophages by promoting lysosomal degradation of TLR4. Blood. 2007;110(3):962–71.

10. da Silva Correia J, Ulevitch RJ. MD-2 and TLR4 N-linked glycosylations are important for a functional lipopolysaccharide receptor. J Biol Chem. 2002;277(3):1845–54.

11. Weber AN, Morse MA, Gay NJ. Four N-linked glycosylation sites in human toll-like receptor 2 cooperate to direct efficient biosynthesis and secretion. J Biol Chem. 2004;279(33):34589–94.

12. de Bouteiller O, Merck E, Hasan UA, Hubac S, Benguigui B, Trinchieri G, et al. Recognition of double-stranded RNA by human toll-like receptor 3 and downstream receptor signaling requires multimerization and an acidic pH. J Biol Chem. 2005;280(46):38133–45.

13. Kirchhausen T. Clathrin. Annu Rev Biochem. 2000;69:699–727.

14. Bonifacino JS, Glick BS. The mechanisms of vesicle budding and fusion. Cell. 2004;116(2):153–66.

15. Lee MC, Miller EA, Goldberg J, Orci L, Schekman R. Bi-directional protein transport between the ER and Golgi. Annu Rev Cell Dev Biol. 2004;20:87–123.

16. Bannykh SI, Rowe T, Balch WE. The organization of endoplasmic reticulum export complexes. J Cell Biol. 1996;135(1):19–35.

17. Hammond AT, Glick BS. Dynamics of transitional endoplasmic reticulum sites in vertebrate cells. Mol Biol Cell. 2000;11(9):3013–30.

18. Rossanese OW, Soderholm J, Bevis BJ, Sears IB, O’Connor J, Williamson EK, et al. Golgi structure correlates with transitional endoplasmic reticulum organization in Pichia pastoris and Saccharomyces cerevisiae. J Cell Biol. 1999;145(1):69–81.

19. Barlowe C, Orci L, Yeung T, Hosobuchi M, Hamamoto S, Salama N, et al. COPII: a membrane coat formed by Sec proteins that drive vesicle budding from the endoplasmic reticulum. Cell. 1994;77(6):895–907.

20. Matsuoka K, Orci L, Amherdt M, Bednarek SY, Hamamoto S, Schekman R, et al. COPII-coated vesicle formation reconstituted with purified coat proteins and chemically defined liposomes. Cell. 1998;93(2):263–75.

21. Antonny B, Beraud-Dufour S, Chardin P, Chabre M. N-terminal hydrophobic residues of the G-protein ADP-ribosylation factor-1 insert into membrane phospholipids upon GDP to GTP exchange. Biochemistry. 1997;36(15):4675–84.

22. Fath S, Mancias JD, Bi X, Goldberg J. Structure and organization of coat proteins in the COPII cage. Cell. 2007;129(7):1325–36.

23. Matsuoka K, Schekman R, Orci L, Heuser JE. Surface structure of the COPII-coated vesicle. Proc Natl Acad Sci U S A. 2001;98(24):13705–9.

24. Stagg SM, LaPointe P, Razvi A, Gurkan C, Potter CS, Carragher B, et al. Structural basis for cargo regulation of COPII coat assembly. Cell. 2008;134(3):474–84.

25. Gurkan C, Stagg SM, Lapointe P, Balch WE. The COPII cage: unifying principles of vesicle coat assembly. Nat Rev Mol Cell Biol. 2006;7(10):727–38.

26. Schimmoller F, Singer-Kruger B, Schroder S, Kruger U, Barlowe C, Riezman H. The absence of Emp24p, a component of ER-derived COPII-coated vesicles, causes a defect in transport of selected proteins to the Golgi. The EMBO journal. 1995;14(7):1329–39.

27. Bonnon C, Wendeler MW, Paccaud JP, Hauri HP. Selective export of human GPI-anchored proteins from the endoplasmic reticulum. J Cell Sci. 2010;123(Pt 10):1705–15.

28. Contreras FX, Ernst AM, Haberkant P, Bjorkholm P, Lindahl E, Gonen B, et al. Molecular recognition of a single sphingolipid species by a protein’s transmembrane domain. Nature. 2012;481(7382):525–9.

29. Fullekrug J, Suganuma T, Tang BL, Hong W, Storrie B, Nilsson T. Localization and recycling of gp27 (hp24gamma3): complex formation with other p24 family members. Molecular biology of the cell. 1999;10(6):1939–55.

30. Marzioch M, Henthorn DC, Herrmann JM, Wilson R, Thomas DY, Bergeron JJ, et al. Erp1p and Erp2p, partners for Emp24p and Erv25p in a yeast p24 complex. Mol Biol Cell. 1999;10(6):1923–38.

31. Jenne N, Frey K, Brugger B, Wieland FT. Oligomeric state and stoichiometry of p24 proteins in the early secretory pathway. The Journal of biological chemistry. 2002;277(48):46504–11.

32. Nagae M, Hirata T, Morita-Matsumoto K, Theiler R, Fujita M, Kinoshita T, et al. 3D Structure and Interaction of p24beta and p24delta Golgi Dynamics Domains: Implication for p24 Complex Formation and Cargo Transport. J Mol Biol. 2016;428(20):4087–99.

33. Manzano-Lopez J, Perez-Linero AM, Aguilera-Romero A, Martin ME, Okano T, Silva DV, et al. COPII coat composition is actively regulated by luminal cargo maturation. Curr Biol. 2015;25(2):152–62.

34. Braakman I, Bulleid NJ. Protein folding and modification in the mammalian endoplasmic reticulum. Annu Rev Biochem. 2011;80:71–99.

35. Breitling J, Aebi M. N-linked protein glycosylation in the endoplasmic reticulum. Cold Spring Harb Perspect Biol. 2013;5(8):a013359.

36. Fujita M, Watanabe R, Jaensch N, Romanova-Michaelides M, Satoh T, Kato M, et al. Sorting of GPI-anchored proteins into ER exit sites by p24 proteins is dependent on remodeled GPI. The Journal of cell biology. 2011;194(1):61–75.

37. Takida S, Maeda Y, Kinoshita T. Mammalian GPI-anchored proteins require p24 proteins for their efficient transport from the ER to the plasma membrane. Biochem J. 2008;409(2):555–62.

38. Astuti GDN, van den Born LI, Khan MI, Hamel CP, Bocquet B, Manes G, et al. Identification of Inherited Retinal Disease-Associated Genetic Variants in 11 Candidate Genes. Genes (Basel). 2018;9(1).

39. Denzel A, Otto F, Girod A, Pepperkok R, Watson R, Rosewell I, et al. The p24 family member p23 is required for early embryonic development. Current biology : CB. 2000;10(1):55–8.

40. Blum R, Pfeiffer F, Feick P, Nastainczyk W, Kohler B, Schafer KH, et al. Intracellular localization and in vivo trafficking of p24A and p23. J Cell Sci. 1999;112 (Pt 4):537–48.

41. Emery G, Rojo M, Gruenberg J. Coupled transport of p24 family members. Journal of cell science. 2000;113 (Pt 13):2507–16.

42. Jerome-Majewska LA, Achkar T, Luo L, Lupu F, Lacy E. The trafficking protein Tmed2/p24beta(1) is required for morphogenesis of the mouse embryo and placenta. Developmental biology. 2010;341(1):154–66.

43. Vavassori S, Cortini M, Masui S, Sannino S, Anelli T, Caserta IR, et al. A pH-regulated quality control cycle for surveillance of secretory protein assembly. Mol Cell. 2013;50(6):783–92.

44. Appenzeller-Herzog C, Roche AC, Nufer O, Hauri HP. pH-induced conversion of the transport lectin ERGIC-53 triggers glycoprotein release. J Biol Chem. 2004;279(13):12943–50.

45. Sun MS, Zhang J, Jiang LQ, Pan YX, Tan JY, Yu F, et al. TMED2 Potentiates Cellular IFN Responses to DNA Viruses by Reinforcing MITA Dimerization and Facilitating Its Trafficking. Cell Rep. 2018;25(11):3086–98 e3.

